# Effect of Thrombospondin-4 on Pro-inflammatory Phenotype Differentiation and Apoptosis in Macrophages

**DOI:** 10.1101/633537

**Authors:** Mohammed Tanjimur Rahman, Santoshi Muppala, Jiahui Wu, Irene Krukovets, Dmitry Solovjev, Dmitriy Verbovetskiy, Edward F. Plow, Olga Stenina-Adognravi

**Affiliations:** Department of Cardiovascular & Metabolic Sciences, Cleveland Clinic, Cleveland, Ohio

**Keywords:** thrombospondin-4, macrophage, inflammation

## Abstract

Thrombospondin-4 (TSP4) attracted a lot of attention recently as a result of new functions identified for this matricellular protein in cardiovascular, muscular, and nervous systems. We have previously reported that TSP4 promotes local vascular inflammation in mouse atherosclerosis model. A common variant of TSP4, P387-TSP4, was associated with increased cardiovascular disease risk in human population studies. In a mouse atherosclerosis model, TSP4 had profound effect on accumulation of macrophages in lesions, which prompted us to examine its effects on macrophages, more in detail in this report.

We examined the effects of A387-TSP-4 and P387-TSP-4 on mouse macrophages in cell culture and *in vivo* in the model of LPS-induced peritonitis. In tissues and in cell culture, TSP4 expression was associated with inflammation: TSP4 expression was upregulated in peritoneal tissues in LPS-induced peritonitis, and pro-inflammatory signals, INFγ, GM-CSF, and LPS, induced TSP4 expression in macrophages *in vivo* and in cell culture. Deficiency in TSP-4 in macrophages from *Thbs4*^−/−^ mice reduced the expression of pro-inflammatory macrophage markers, suggesting that TSP-4 facilitates macrophage differentiation into pro-inflammatory phenotype. Expression of TSP4, especially more active P387-TSP4, was associated with higher cellular apoptosis. Cultured macrophages displayed increased adhesion to TSP4 and reduced migration in presence of TSP4, and these responses were further increased with P387 variant.

We concluded that TSP4 expression in tissue macrophages and in cultured macrophages increases their accumulation in tissues during the acute inflammatory process and supports macrophage differentiation into a pro-inflammatory phenotype. In a model of acute inflammation, TSP4 supports pro-inflammatory macrophage apoptosis, a response that is closely related to their pro-inflammatory activity and release of pro-inflammatory signals. P387-TSP4 was found to be more active form of TSP4 in all examined functions.

## Introduction

Thrombospondin-4 (TSP4) is a matricellular protein, one of the five members of thrombospondin family [1,2]. TSP4 expression is low in adult tissues but dramatically increases during tissue remodeling and regeneration. High levels of TSP4 have been detected in remodeling and failing hearts [3–7], several cancers [8–13], and atherosclerotic lesions [14]. Recently, new functions have been ascribed to TSP4 in cardiovascular system [5–7,14–25], cancer [8–12,24–29], and nervous system [30–34], but effects of TSP4 on cellular responses and the regulation of TSP4 expression remain poorly understood. For example, with an exception of TGF-beta [25], stimuli inducing TSP4 expression in tissues remain unknown. Although several cell surface receptors were shown to mediate TSP4 effects [14,24,30,35], signaling and regulation of intracellular processes by TSP4 have received only limited attention.

Tissue remodeling, inflammation, and angiogenesis often occur simultaneously and are regulated by the same stimuli. We recently reported that TSP4 promotes angiogenesis [24,25] and vascular inflammation in a mouse atherosclerosis model [14]. These observations suggested that TSP4 may play an important role in regulation of inflammation and inflammatory cell responses. In the present study, we describe effects of TSP4 and its common variant, P387-TSP4, on macrophages. The P387-TSP-4 variant is associated with cardiovascular disease and myocardial infarction [15–22] and is more active than WT A387-TSP4 in mediating cellular effects [24,35–37]. TSP4 production by blood cells has not been reported to date. Here, we report several stimuli that induce or reduce TSP4 production in macrophages. The effect of TSP4 on macrophages has been examined in isolated macrophages and in LPS-induced peritonitis, an *in vivo* model of acute inflammation.

Macrophages play multiple roles in both the tissue homeostasis and inflammation, e.g., promote the initiation, progression, and healing of tissue injury; facilitate tissue remodeling associated with various pathologies; and participate in initiation and resolution of inflammation in tissues [38,39]. Macrophages are a heterogeneous cell types with high plasticity and display a variety of phenotypes regulated by external stimuli, including the extracellular matrix [40]. A continuum of phenotypes exists that can be pragmatically divided into pro-inflammatory and tissue repair macrophages. Phenotypic polarization of macrophages is regulated by pro-inflammatory stimuli that include Toll-like receptor (TLR) ligands and IFNγ, which induce pro-inflammatory phenotypes, and by anti-inflammatory stimuli such as IL4/IL13, immune complexes, the anti-inflammatory cytokines IL-10, and transforming growth factor-β that induce tissue repair phenotypes [41–44]. Although the effects of microenvironment, including ECM proteins and matricellular proteins specifically, on phenotypic differentiation of macrophages are undeniable, they are still poorly understood. Here, we have uncovered previously unrecognized cellular mechanisms for regulation of phenotypes and functions of macrophages by TSP4 and the P387-TSP4 variant.

## Methods and Materials

### Animals

Mice were of the C57BL/6 background. Both genders were used (there were no gender-specific differences). *Thbs4*^−/−^ mice were described previously [5,14,24,25,45]. Control wild-type (WT) mice were from the same mouse colony as *Thbs4*^−/−^ mice or P387-TSP4-KI mice. For experiments exclusively using WT mice, WT C57BL/6 mice were purchased from the Jackson Laboratories. Animals were housed in the AAALAC-approved animal facilities of the Cleveland Clinic. Animal procedures were approved by y the Institutional Animal Care and Use Committee of the Cleveland Clinic and were in agreement with the NIH Guide for Animal Use.

### LPS Induced peritonitis

Mice were treated with intraperitoneal (IP) injection of Lipopolysaccharide (LPS, 0.5 μg/g, ThermoFisher) to induce peritonitis 72 hours prior to isolation of peritoneal cells by lavage and resection of visceral peritoneum. Mouse peritoneal macrophages (MPM) were used directly for cell adhesion and migration experiments. mRNA was isolated from MPM and peritoneal tissue. Immunofluorescence was also done on peritoneal tissues. IP injection with PBS was used for the control groups of animals.

### Isolation and culture of primary macrophages

Bone-marrow-derived macrophages (BMDM) were isolated from hind leg tibia and femur of WT, *Thbs4*^−/−^ or P387-TSP-4 KI mice. The bone marrow cells were collected and differentiated in to BMDM using 30 ng/mL M-CSF for 7 days. These cells were treated with 50 ng/mL GM-CSF, 50 ng/mL M-CSF, 40 ng/mL IL-4, 2000 IU/mL IFNγ, 0.5 μg/mL LPS. Following stimulations, cells were analyzed by western blot and quantitative RT-PCR for specific protein/gene expression patterns.

### Cell Culture and Cell Lines

Mouse macrophage cell line RAW 264.7 was purchased from ATCC (Virginia, USA). DMEM media (RAW 264.7) and grown in Eagle’s Minimum Essential Medium (WBC264-9C) supplemented with 10% fetal bovine serum (FBS), sodium bi-carbonate (1.5gm/L), 100 U/mL penicillin, and 100 µg/mL streptomycin at 37 °C with 5% CO2 atmosphere according to ATCC recommendations. Cells were treated with 20 ng/mL GM-CSF (#NBP2-35066, Novus Biologicals), 20 ng/mL M-CSF (**#**14-8983-62, ThermoFisher), 40 ng/mL IL-4 (#574306, BioLegend), 1000 IU/mL IFNγ (#NBP2-35071, Novus Biologicals), 0.5 μg/mL lLipopolysaccharide (#00-4976-93, ThermoFisher), 25 μM cycloheximide (#0970, R&D Systems) dissolved in serum free DMEM. All RAW 264.7 cell experiments were done with cells between passage 4 and passage 8.

Cells were exposed to a brief period of serum starvation in serum-free DMEM for 3 hours followed by stimulation with pro-inflammatory stimuli (LPS, GM-CSF, IFNγ), anti-inflammatory stimuli (M-CSF, IL-4) and cycloheximide (CHX) at above-mentioned doses for stated durations.

### Western Blot Analysis

Cells were harvested and lysed on ice using RIPA Lysis and Extraction Buffer (# 89901, Thermo Scientific) supplemented with Halt™ protease and phosphatase inhibitor cocktail (#78442, Thermo Scientific). Cell lysates were boiled with 2-Mercaptoethanol (#21985-023, LifeTechnologies) activated Laemmli Sample Buffer at 100 °C for 5 minutes. 50 mg of total protein was loaded in each well of 8% a Bis-Tris gel, run and transferred to PVDF blotting membrane using semi dry transfer module. After blocking with 5% skim milk in TBST for an hour at room temperature, membranes were probed overnight at 4 °C with primary antibodies [Anti TSP4 primary monoclonal Ab (#sc-390734, Santa Cruz Biotechnology) at 1:100 dilution; Anti β-Actin Ab (#A5316, Sigma-Aldrich) at 1:10000 dilution] followed by respective HRP anti-mouse antibody incubation and developed with Pierce® ECL Western Blotting Substrate on X-ray films

### Quantitative real-time RT-PCR analysis

Total RNA was extracted using TRIzol Reagent (#15596026, ThermoFisher) followed by cDNA synthesis using SuperScript™ First-Strand Synthesis System (#11904018, ThermoFisher). The TaqMan® probe for *Thbs4* (Assay ID# Mm03003598_s1, ThermoFisher), *CD68* (Assay ID# Mm03047343_m1, ThermoFisher), *CD38* (Assay ID# Mm01220906_m1, ThermoFisher), *Egr2* (Assay ID# Mm00456650_m1, ThermoFisher), *Nos2* (Assay ID# Mm00440502_m1, ThermoFisher), *Arg1* (Assay ID# Mm00475988_m1, ThermoFisher) were used for detecting respective genes expressions with Stx5a (Assay ID# Mm00502335_m1, ThermoFisher)) and *ACTB* (Assay ID# Mm02619580_g1, ThermoFisher) were used as housekeeping gene controls.

### Macrophage differentiation and survival assays

Cells were seeded in 24-well plates at 300,000 cells/well in M1 and M2 differentiation media (#C-28055, #C-28056, Promo Cell). Initial attachment of cells was determined after 3 hours using CyQUANT live cell quantification kits. Cells were kept into M1 and M2 differentiation media for 5 days followed by total live cell quantification using CyQUANT reagent. Each groups of live cell quantification after 5-day time-points were normalized to their respective 3 hours initial attachment quantification and compared to assess cell survival of macrophages from WT or *Thbs4*^−/−^ or P387-TSP4-KI mice.

### BMDM Apoptosis Assay

BMDM isolated from WT, *Thbs4*^−/−^ or P387-TSP-4-KI mice were differentiated with 50 ng/mL GM-CSF for 7 days. BMDM were treated with 0.5 μg/mL LPS in serum free media for 24 hours followed by qRT-PCR for *Bax, Bcl2* and *Caspase 3* (Casp3) expression.

For apoptosis assays, BMDM were seeded into 96-well plates (20,000 cells/well) and treated with 0.5 μg/mL LPS for 48 hours. Apoptosis was measured using Cell Meter™ No-Wash Live Cell Caspase 3/7 Activity Assay Kits (#20250, AAT Bioquest). Caspase 3/7 activity data were normalized to total number of live cells (determined by CyQUANT kit) in each of the respective experimental groups.

### In vitro adhesion assays

Adhesion assays were performed as previously described [14,24,25,35,37]. MPM were isolated from WT, *Thbs4*^−/−^ and P387-TSP4-KI mice 72 hours after treatment with 0.25μg/g LPS, and 5 × 10^3^ cells were added for 1 hour at 37 °C to wells of 24-well plates (Corning) pre-coated with fibronectin (Sigma-Aldrich) or without coating.

### In vitro migration assays

MPM were isolated from LPS treated WT, *Thbs4*^−/−^ and P387-TSP4-KI mice, and 0.2 × 10^6^ MPM were resuspended in the serum-free DMEM and transferred into the trans-well chambers (Corning, Corning, NY, USA). FBS (20%) was used as a chemo-attractant in the bottom chambers. The cells were incubated at 37 °C for 4 hours, the medium was aspirated, and attached cells were removed from the surface of the upper chamber using Q-tips. The plates were frozen at −80 °C for 3 hours, and DNA from remaining cells was quantified using CyQUANT reagent (Invitrogen, Carlsbad, CA, USA).

### Immunohistochemistry, Immunofluorescence, confocal imaging and quantification of macrophage markers

Sections of mice peritoneum were stained with primary rat anti-mouse CD68 antibody (MCA1957B, Bio-Rad), rabbit anti-mouse CD38 antibody (bs-0979R, Bioss Antibodies), Rabbit anti-mouse Egr2 (ab90518, Abcam) antibody using Vecta Stain ABC Kit. Visualization after staining with the antibodies was performed using a high-resolution slide scanner, Leica Aperio AT2 slide scanner (Leica Microsystems, GmbH, Wetzlar, Germany) was used to scan images of whole slides at 20x magnification and quantified to determine the percentage of the stained area using Adobe photoshop CS6 (Media Cybernetics). The person performing quantification was blinded to the assignment of animals between groups. For immunofluorescence, rat anti-mouse CD68 (MCA1957B, Bio-Rad), goat anti-human TSP4 (AF2390, R&D systems), were used with corresponding secondary antibodies (1:1000). Secondary antibodies were: anti-goat NL557 conjugated donkey IgG (NL001 R&D systems) and goat polyclonal antibody to rat IgG Alexa Fluor 488 (ab150161, abcam). Images were taken at a high-resolution confocal microscope (Leica DM 2500) at 63× magnification. All sections with primary antibodies were incubated for 2 hours at 4°C followed by incubating sections in secondary antibodies for 45 min at 4°C.

### Statistical analysis

Group size was calculated based on the previous data obtained in mouse models [5,14,24,45]. Analyses of the data were performed using Sigma Plot Software (Systat Software, San Jose, CA, USA): Student’s t-test and one-way ANOVA were used to determine the significance of parametric data, and Wilcoxon rank sum test was used for nonparametric data. The significance level was set at p=0.05. The data are presented as mean ± SEM, number of biological repeats are listed in each legend.

## Results

### LPS and pro-inflammatory cytokines induce TSP-4 expression in macrophages

LPS induced TSP4 in a time-dependent manner in RAW264.7 and BMDM (Fig.1A). Most TSP-4 was intracellular in both macrophage types (Fig. 1B). In both cell types, both the protein levels and mRNA levels were increased in response to LPS stimulation of cells (Fig. 1A and C). The decreases in protein levels in cells treated with cycloheximide were comparable to untreated cells, also consistent with the transcriptional mechanism or increased RNA stability.

**Figure 1.**
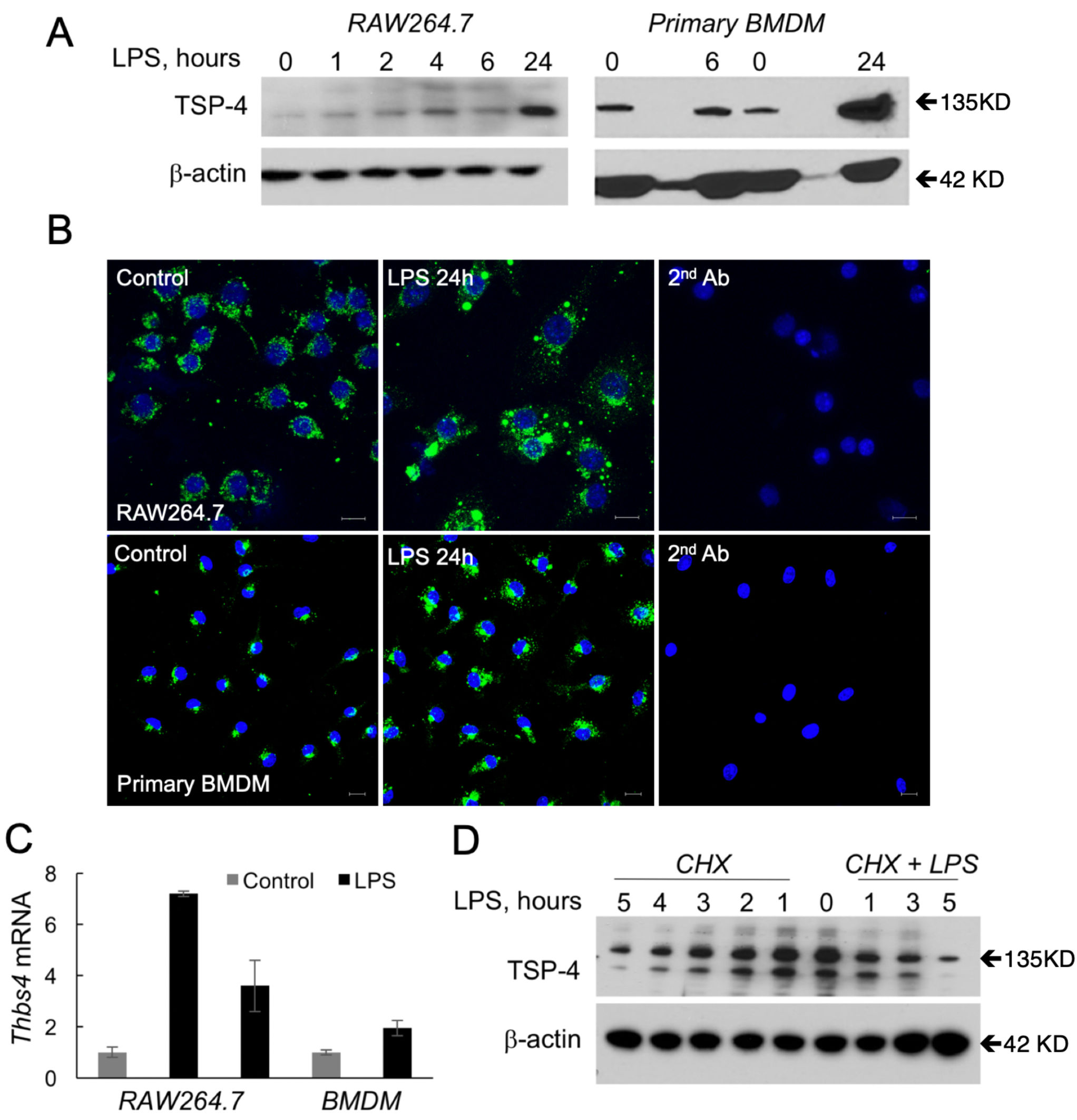
LPS induces TSP-4 in cultured macrophages. **A:** Cultured macrophage-like RAW264.7 and cells and mouse bone-marrow derived macrophages (BMDM) were treated with LPS (0.5μg/ml for 1 – 24 hours); Western blotting with anti-TSP-4 Ab. **B:** Cultured RAW264.7 (upper panels) and BMDM (lower panels) treated with LPS for 24 hours were stained with anti-TSP-4 Ab; green = TSP-4, blue = nuclei, DAPI. Scale bar is 10µm. **C:** RAW 264.7 and BMDM cells were treated with LPS (0.5μg/mL) for 24 hours and qRT-PCR was done (control = PBS treated); **D:** RAW 264.7 cells were treated with CHX (25μM) for 1-5 hours and CHX (25μM) + LPS (0.5μg/mL) for 1-3 hours followed by Western blot detection of TSP-4 (β-actin as loading controls).

We examined the effects of pro-inflammatory (LPS, IFNγ, and GM-CSF) and anti-inflammatory stimuli (M-CSF and IL-4) on the levels of TSP-4 protein and mRNA in RAW 264.7 cells (Fig. 2) and in BMDM (Fig.3). BMDM were isolated from mice and cultured in macrophage differentiation media for 7 days. The purity was assessed by flow cytometry using anti-CD11b and F4/80 staining and was 93-98% (Suppl. Fig.1). Western blot analysis showed a gradual increase in TSP-4 protein levels with pro-inflammatory stimuli LPS, IFNγ and GM-CSF. Conversely, TSP-4 protein levels reduced with anti-inflammatory cytokine IL-4 and M-CSF stimulation (Fig. 2A and B; Fig.3A). Average TSP-4 protein amount was quantified from three independent experiments and normalized to β Actin expression as a loading control. Anti-inflammatory stimuli caused significant (\**p*<0.05) reduction in TSP-4 protein levels (Fig. 2A and B, Fig.3A and B).

**Figure 2.**
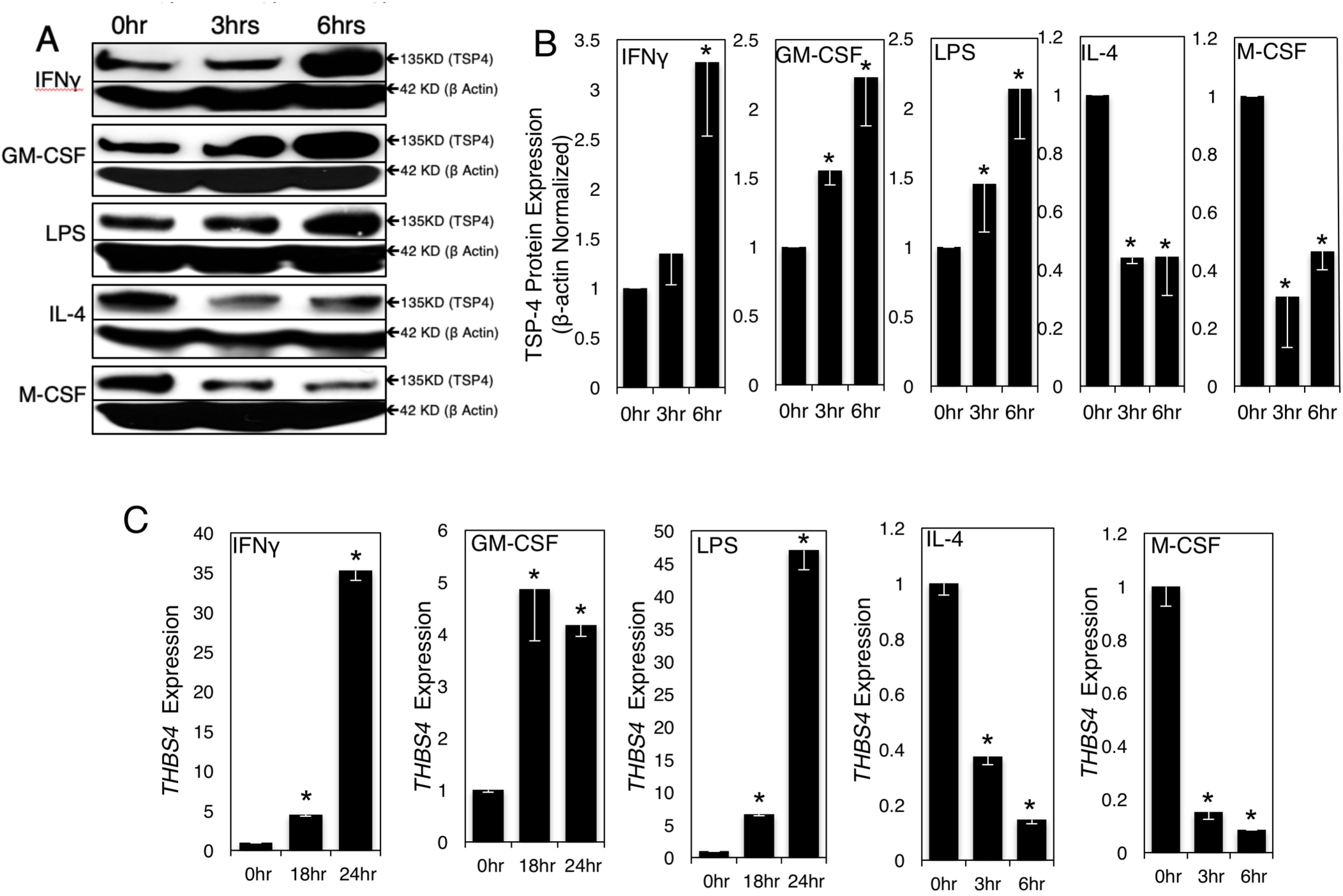
Pro-inflammatory stimuli upregulate TSP-4 expression in cultured macrophage-like cell line RAW264.7. Cultured RAW264.7 were treated with pro-inflammatory (1000 IU/ml INF-γ, 20 ng/ml GM-CSF, 0.5 μg/ml LPS) and anti-inflammatory (40 ng/ml IL-4, 20 ng/ml M-CSF) stimuli for indicated times. **A:** TSP-4 protein was detected in Western blotting with anti-TSP-4 Ab, representative images. **B:** Protein levels were quantified by densitometry; n = 3; p <0.05. **C:** *Thbs4* expression was measured by QRT-PCR; n = 3; *p <0.05.

*THBS4 mRNA levels* was measured by quantitative RT-PCR (Fig.2C and 3C). There was significant (\**p*<0.05) increase in *THBS4* expression upon stimulation with pro-inflammatory stimuli and a decrease in *THBS4* expression with anti-inflammatory stimuli.

Expression of markers of both pro-inflammatory (CD38 and Nos2, Suppl. Fig. 2) and tissue-repair (Egr2 and Arg1, Suppl. Fig. 3) macrophages was measured in RAW 264.7 cells and BMDM treated. Following stimulation with IFNγ (1000IU/mL), GM-CSF (20ng/mL), LPS (0.5μg/mL), M-CSF (20ng/mL) or IL-4 (40ng/mL), increased TSP-4 levels were detected and correlated with increased CD38 and Nos2 expression and a decrease in Egr2 and Arg1 expression. Decrease in TSP-4 levels was associated with decreased CD38 and Nos2 expression and increased Egr2 and Arg1 expression.

**Figure 3.**
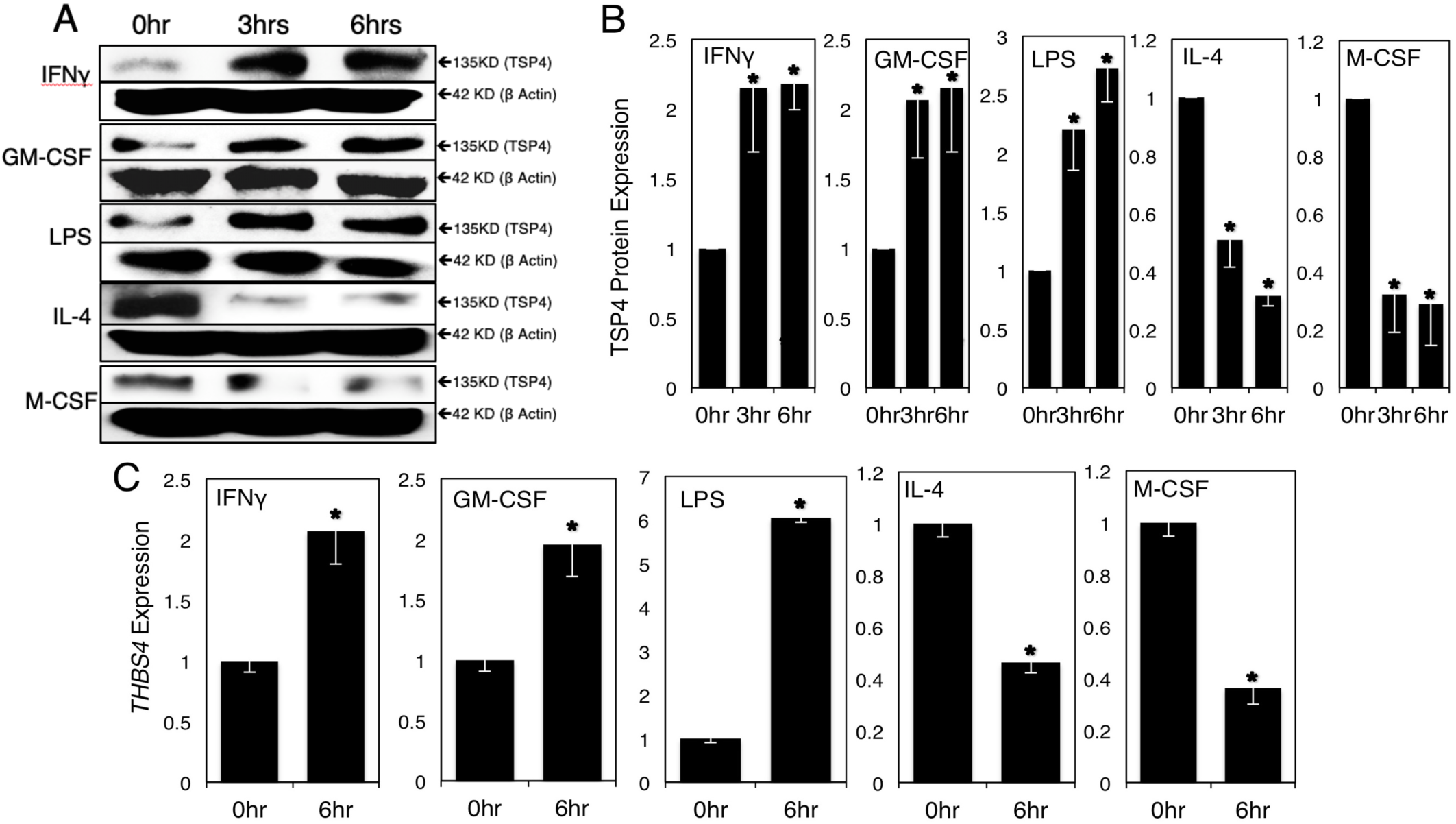
Pro-inflammatory stimuli upregulate TSP-4 expression in cultured mouse bone-marrow-derived macrophages. Cultured BMDM were treated with pro-inflammatory (1000 IU/ml INF-γ, 20 ng/ml GM-CSF, 0.5 μg/ml LPS) and anti-inflammatory (40 ng/ml IL-4, 20 ng/ml M-CSF) stimuli for indicated times. **A:** TSP-4 protein was detected in Western blotting with anti-TSP-4 Ab, representative images. **B:** Protein levels were quantified by densitometry; n = 3; *p <0.05. **C:** *Thbs4* expression was measured by QRT-PCR; n = 3; *p <0.05.

**Figure 4.**
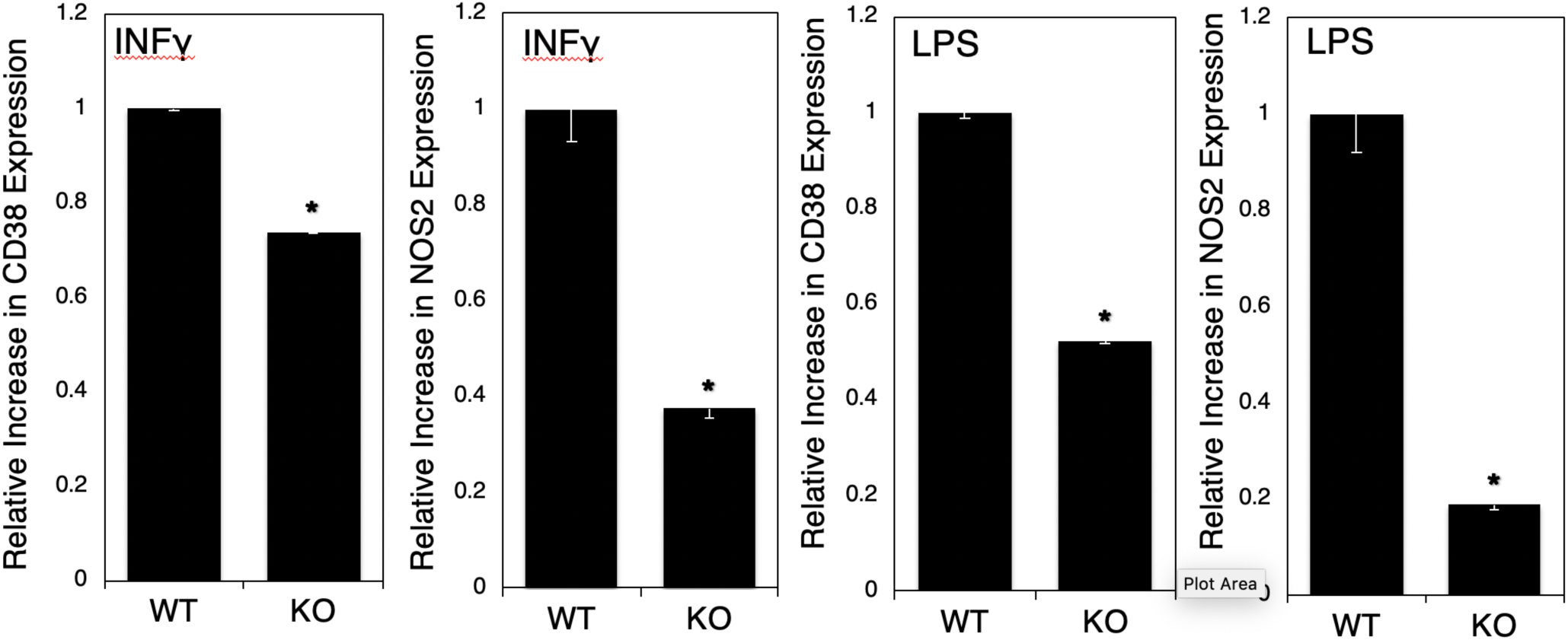
Deletion of Thbs4 in mouse BMDM decreases the levels of pro-inflammatory macrophage markers. BMDM were treated with LPS (0.5 μg/mL) and IFNγ (1000 IU/mL) for 6 hours, and Nos2 and CD38 mRNA expression was measured by QRT-PCR; n = 3; *p <0.05.

### TSP-4 promotes differentiation of macrophages into pro-inflammatory phenotype

To understand whether TSP-4 promotes the differentiation of macrophages into pro-inflammatory phenotype or whether the increased production of TSP-4 results from differentiation but does not affect the process, BMDM from WT and *Thbs4*^−/−^ mice were stimulated with LPS (0.5 μg/mL) and IFNγ (1000 IU/mL), and the levels of mRNA of the markers of pro-inflammatory differentiation of macrophages CD38 and Nos2 were measured by Real-Time PCR (Fig.4). The levels of both markers were significantly decreased in BMDM from *Thbs4*^−/−^ mice, suggesting that TSP-4 promotes the differentiation into proinflammatory phenotype.

### TSP-4 regulates the survival of pro-inflammatory macrophages

Programmed death is a natural fate of pro-inflammatory macrophages and is associated with release of inflammatory signals [46–53]. Knowing that TSP-4 expression promotes pro-inflammatory phenotype of macrophages, we investigated the effect of TSP-4 on cultured macrophage apoptosis.

When BMDM from WT, *Thbs4*^−/−^, and P387-TSP4-KI mice were differentiated in pro-inflammatory (M1) medium or tissue repair differentiation (M2 medium), the survival of cultured pro-inflammatory BMDM but not tissue repair BMDM was increased in the *Thbs4*^−/−^ cells and decreased in P387-TSP4-KI cells (Fig. 5A).

**Figure 5.**
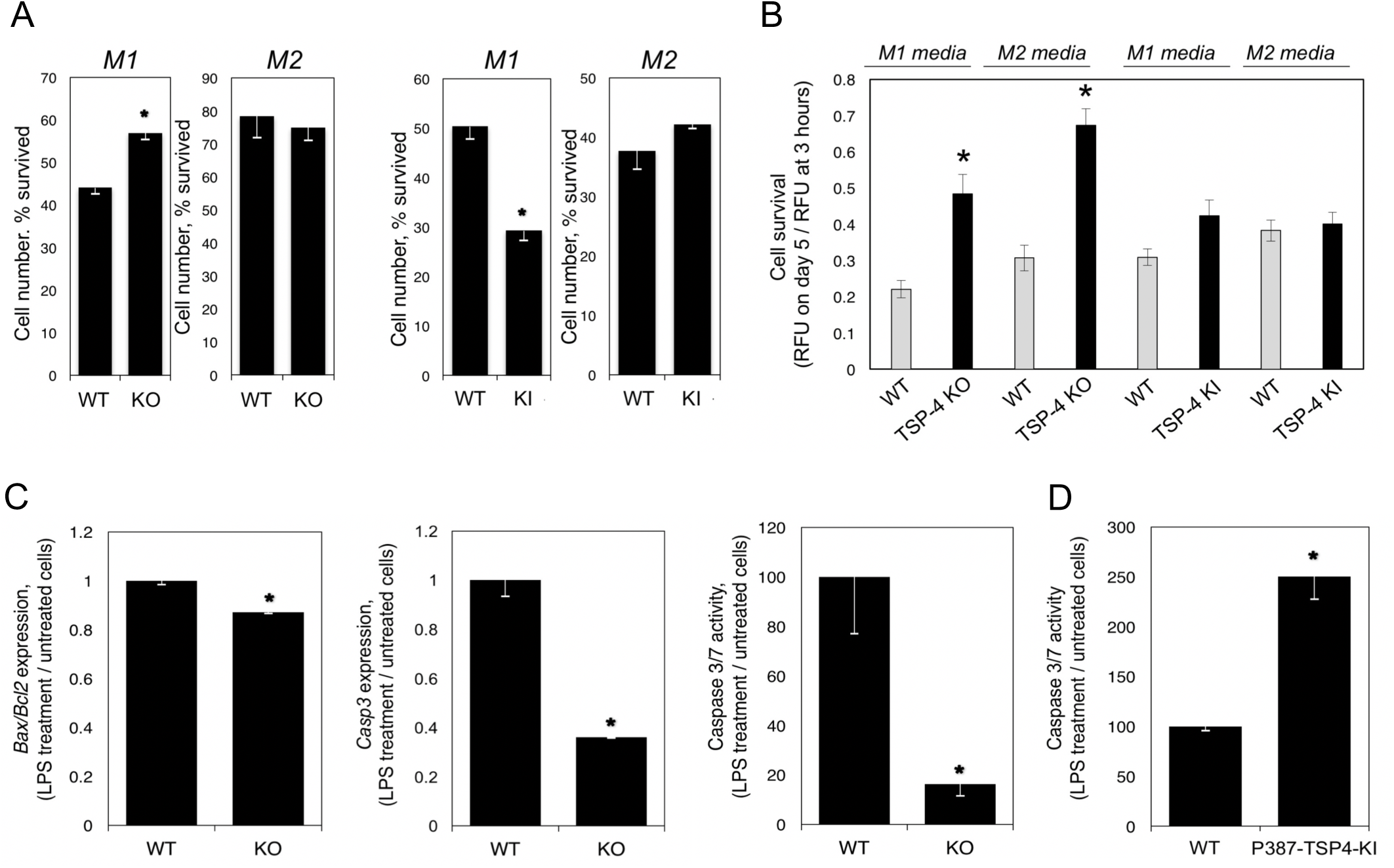
TSP-4 promotes decreases survival and promotes apoptosis in pro-inflammatory macrophages. **A:** Bone-marrow-derived macrophages (BMDM) from WT, KO, and KI mice were differentiated in culture in M1 or M2 differentiation media, and the number of survived cells (number of cells on day 5/number of cells 3 hour after seeding) was measured using the CyQuant cell survival assay kit; n=5; *p<0.05. **B:** Purified mouse blood monocytes were differentiated in M1 or M2 differentiation cell culture media, and the number of survived cells was measured; n=12; *p<0.05. **C:** Bone-marrow-derived macrophages (BMDM) were differentiated in M1 differentiation medium for 7 days and stimulated with LPS for 48 hours. The expression of *Bax, Bcl2*, and *Casp3* and the activity of Caspase 3/7 enzymes were measured in cultured BMDM from WT and *Thbs4*^−/−^ mice; n = 3; *p < 0.05. **D:** The activity of Caspase 3/7 enzymes was measured in cultured BMDM from WT and P387-TSP4-K*I* mice; n = 3; *p < 0.05.

Monocytes were isolated from mouse blood and differentiated in M1 or M2 media. In M1 or M2 media, lack of TSP-4 resulted significantly higher survival of cells, but P387-TSP-4 did not affect cell number (Fig. 5B).

### TSP-4 promotes apoptosis in pro-inflammatory macrophages

Expression of *Bax, Bcl2, Casp3* and the activity of Caspase 3/7 were measured in BMDM differentiated into pro-inflammatory macrophages (Fig. 5C, D). Lower apoptotic activity was detected in *Thbs4*^−/−^ cells as indicated by decreased *Bax/Bcl2* ratio, decreased expression of *Casp3*, and activity of Caspase 3/7 (Fig.5C). In contrast, the activity of Caspase 3/7 was upregulated in response to LPS in P387-TSP4-KI BMDM (Fig.5D). This differential effect is consistent with the higher activity of P387-TSP4 in other cellular responses [24,35–37].

### Macrophages in peritoneal cavity and tissue of mice with LPS-induced peritonitis

To evaluate the expression and the effect of TSP-4 on macrophages in a model of acute inflammation, we induced peritonitis in WT, *Thbs4*^−/−^, and P387-TSP4-KI mice by IP injection of LPS. Mice were sacrificed 72 hours later, and cells (> 90%) macrophages) were collected by peritoneal lavage and quantified (Fig. 6A). Surprisingly, TSP4 deletion in the *Thbs4*^−/−^ mice did not reduce the number of macrophages recovered in a saline lavage (free macrophages) compared to WT mice, but we collected fewer macrophages from the cavity of P387-TSP4-KI mice than from WT mice after LPS-induced peritonitis (Fig. 6A). To find out whether the LPS-induced inflammation affects the expression of TSP-4 in peritoneal tissues and in free macrophages, we assessed the levels of *Thbs4* mRNA (Fig.6B). TSP-4 expression was dramatically decreased in free peritoneal macrophages and significantly increased in peritoneal tissue of WT mice, suggesting that high levels of TSP-4 in tissue and in macrophages may promote their accumulation in the peritoneal tissue and/or prevent their egress into the cavity.

**Figure 6.**
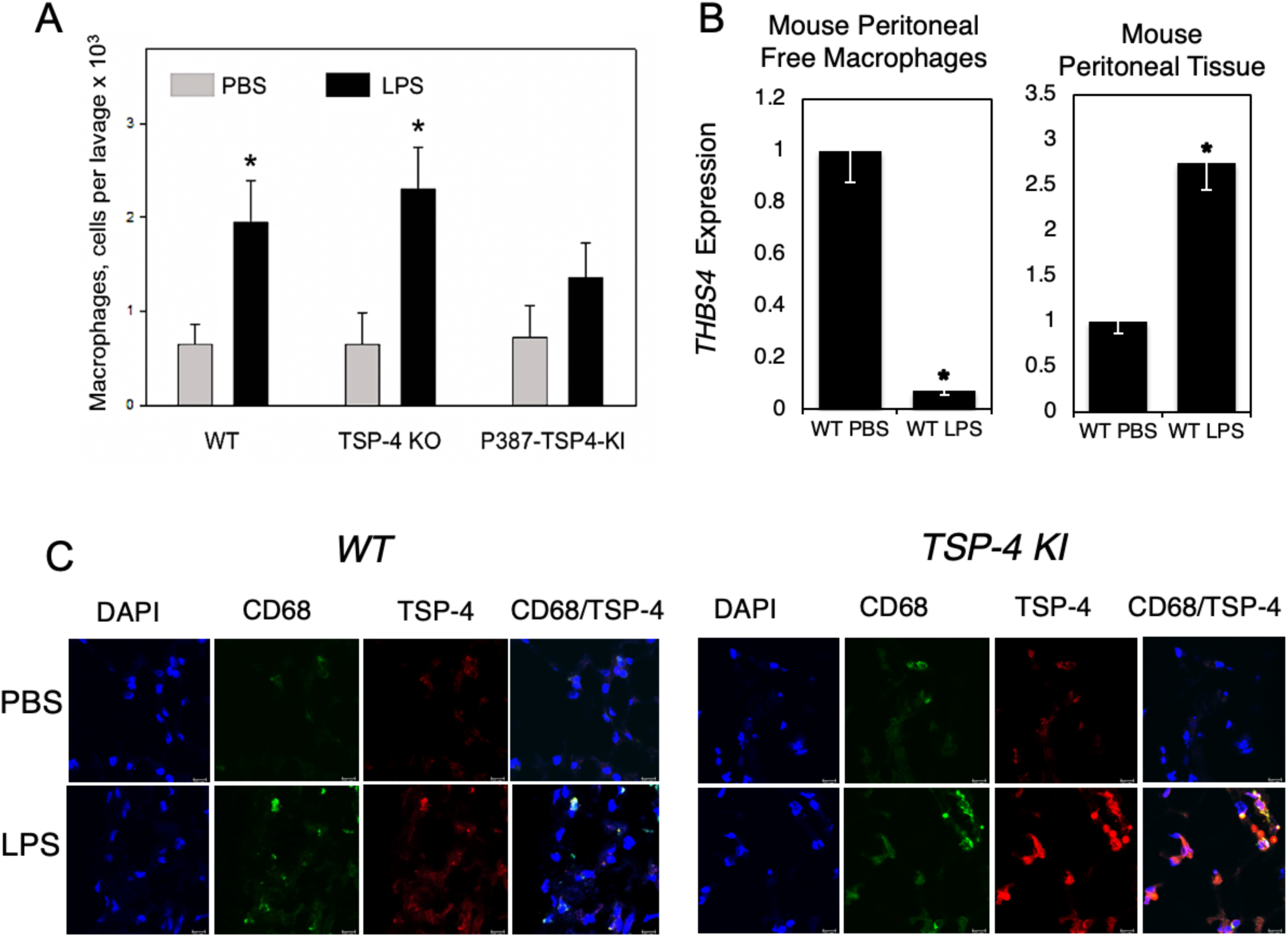
TSP-4 promotes accumulation of macrophages in peritoneal tissue of mice with LPS-induced peritonitis. **A:** Number of macrophages in peritoneal cavity in mice with LPS-induced peritonitis. *p < 0.05, n = 5. **B:** TSP-4 expression in macrophages from the peritoneal cavity lavage (left panel) and in the peritoneal tissue. QRT-PCR, fold increase (RQ) over the values in control mice injected with PBS; n = 3; *p < 0.05. **C:** Macrophages and TSP-4 in peritoneal tissue of WT and P387 TSP-4 KI mice with LPS-induced peritonitis. Immunofluorescence; blue = nuclei (DAPI), green = macrophages (anti-CD68), red = TSP-4 (anti-TSP-4). Scale bar is 10µm.

Immunofluorescence with anti-CD68 and anti-TSP-4 antibodies revealed that TSP-4 protein is associated with macrophages in peritoneal tissue in both WT and P387-TSP4-KI mice (Fig. 6C).

There were no significant changes in monocyte counts in the blood from the two transgenic mice (Suppl. Fig. 4).

### TSP4 increases the number of pro-inflammatory macrophages in peritoneal tissue

Expression of CD68, a marker of macrophages, was upregulated in peritoneal tissues of WT (Fig.7A) and even more dramatically in *Thbs4*^−/−^ (Fig.7B) mice with LPS-induced peritonitis. TSP4 deficiency resulted in higher accumulation of macrophages in peritoneal tissues. In a view of a lack of differences in numbers of macrophages in peritoneal cavity (Fig.6A) and the equal numbers of macrophages in blood (Suppl. Fig.4), this suggested the effect of TSP4 on macrophage survival in tissues, consistent with our observations in cultured macrophages. The expression of the marker of pro-inflammatory macrophages CD38 was upregulated in both WT and *Thbs4*^−/−^mice (Fig.7A and B), but the expression of Egr2, a marker of tissue-repair macrophages, was decreased (Fig. 7A), suggesting that most macrophages retained in the peritoneal tissue are pro-inflammatory. Consistent with the difference between WT and *Thbs4*^−/−^ mice and the P387-TSP4, P387-TSP4 reduced the accumulation of macrophages in peritoneal tissue (Fig.7C), suggesting more profound effect on their survival.

**Figure 7.**
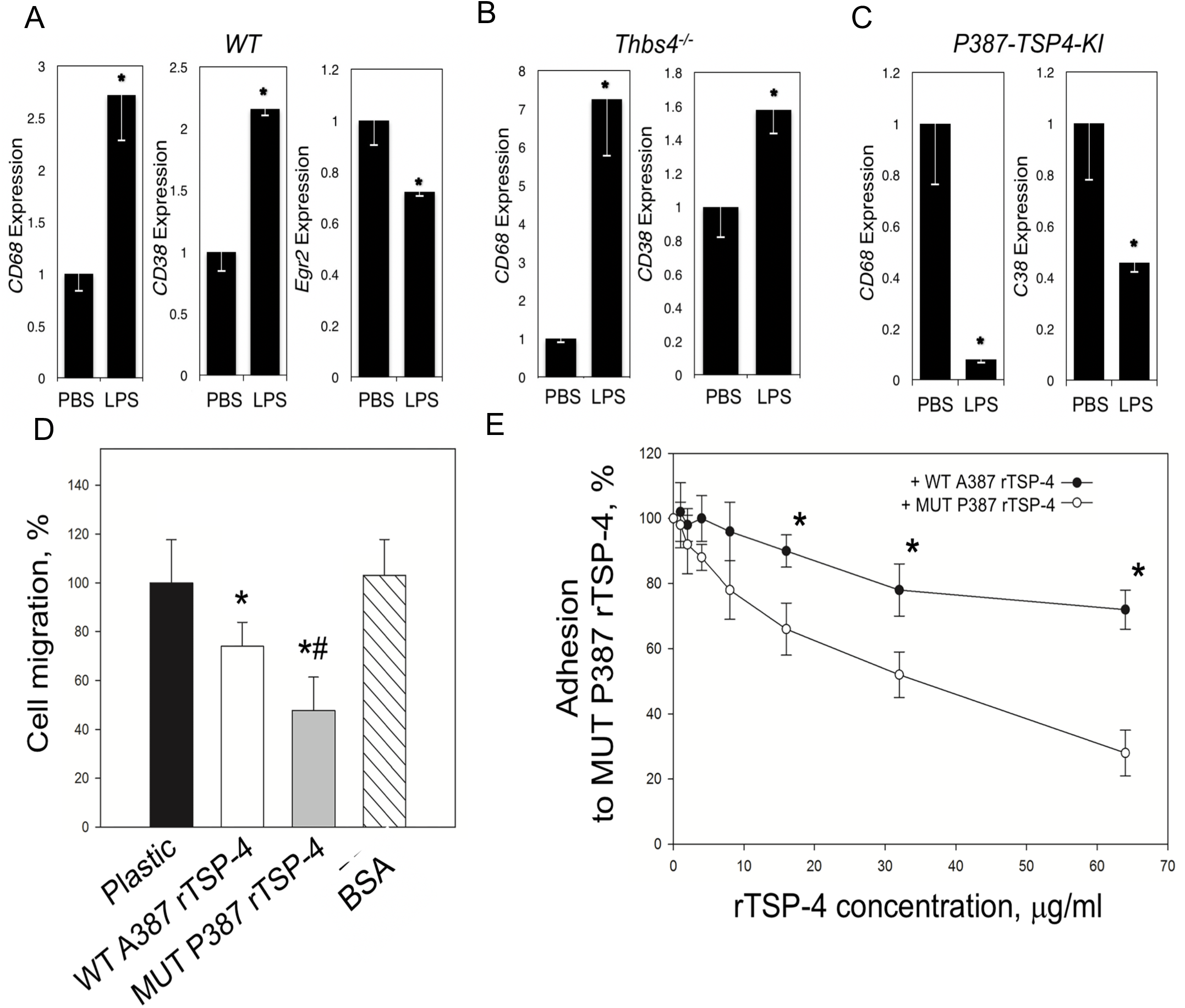
TSP-4 promotes accumulation of pro-inflammatory macrophages in peritoneal tissue of mice with LPS-induced peritonitis. **A:** Expression of CD68, CD38, and Egr2 was measured in peritoneal tissues of WT mice with LPS-induced peritonitis; n=3; *p<0.05. **B:** Expression of CD68 and CD38 was measured in peritoneal tissues of *Thbs4*^−/−^ mice with LPS-induced peritonitis; n=3; *p<0.05. **C:** Expression of CD68 and CD38 was measured in peritoneal tissues of P387-TSP4-KI mice with LPS-induced peritonitis; n=3; *p<0.05. **D:** Migration of RAW264.7 was measured in Boyden chambers (uncoated, coated with A387 or P387 rTSP-4, or BSA); n=3; *p<0.05 compared to plastic and BSA; #p<0.05 compared to WT rTSP-4. **E:** A387 and P387 rTSP-4 was used in adhesion competition assay: plastic was coated with P387 rTSP-4, and RAW264.7 were added in increasing concentrations; n=3; *p<0.05.

### P387 TSP-4 is more active in supporting the adhesion of monocytes and macrophages

We used murine macrophage-like cell line RAW264.7 to assess the effect of A387 TSP-4 and P387 TSP-4 on macrophage adhesion and migration (Fig.7). RAW264.7 cells migrated less on rTSP-4 as a substrate, and P387 TSP-4 further decreased their migration (Fig.7A). RAW264.7 adhered significantly better to P387 TSP-4 (Fig.7B), although both A387 and P387 TSP-4 supported the adhesion of macrophage-like cells.

## Discussion

High expression of TSP4 in remodeling tissues, particularly in heart disease and cancer [5–12,14–29], suggested that this matricellular protein may regulate fibrosis and angiogenesis. The effects of TSP-4 on matrix remodeling has been demonstrated in *Thbs4*^−/−^ mice by us and others [5–7,45], and we recently reported that TSP4 is pro-angiogenic [24,25] in contrast to the prominent role of TSP-1 as an anti-angiogeneic protein [2,54,55]. Tissue remodeling is associated with inflammation, and we reported that TSP4 deficiency results in reduced inflammation in atherosclerotic lesions in *ApoE*^−/−^ mice: the number of macrophages in the atherosclerotic lesion and the local vascular inflammation were reduced in *Thbs4*^−/−^*/ApoE*^−/−^ mice [14]. To investigate the roles of TSP4 and its variants in inflammation, we examined the effects of TSP-4 on cultured macrophages and on macrophages in an LPS-induced mouse peritonitis model.

Several agents that are known to promote pro-inflammatory differentiation of macrophages [LPS, GM-CSF, and INFγ [56]] or tissue-repair differentiation [IL-4 [56] and M-CSF [57]] were used to stimulate cultured RAW264.7 or BMDM. LPS that was also used to induce peritonitis increased TSP-4 production by macrophages in a time-dependent manner. mRNA of *Thbs4* was also upregulated, and production of the protein was efficiently blocked by cycloheximide, suggesting transcriptional regulation or reduced RNA stability. The newly synthesized protein accumulated in vesicle-like structures inside the cells and was not efficiently secreted into the medium or matrix (not shown). Intra-cellular functions have been reported for TSPs [7]. Finding of increased amounts of intracellular TSP4 suggests that such intracellular functions of TSP4 may be activated by the treatment or that macrophages accumulate protein in vesicles that may be released in specific tissues where an inflammation response has been triggered.

All pro-inflammatory signals that resulted in pro-inflammatory differentiation with increased expression of CD38 and Nos2 (markers of pro-inflammatory macrophages) and decreased expression of Egr-2 and Arg1 (markers of tissue repair macrophages) also increased TSP-4 expression and production. To the contrary, anti-inflammatory signals promoting the tissue repair differentiation of macrophages (decreased CD38 and Nos2 and increased Egr-2 and Arg1) decreased the expression of *Thbs4* and the production of the TSP4 protein. Thus, our results suggested a novel role for TSP-4 in inflammation: support of pro-inflammatory functions of macrophages. When BMDM from *Thbs4*^−/−^ mice were stimulated with pro-inflammatory signals, the levels of markers of pro-inflammatory macrophages were decreased, demonstrating that TSP-4 actively promotes macrophage differentiation into pro-inflammatory phenotype.

We analyzed cultured primary BMDM and blood monocytes and found that survival of both cell types was higher in *Thbs4*^−/−^ macrophages cultured in M1 medium. P387-TSP4 in macrophages from P387-TSP4KI mice significantly decreased their survival. Knowing that TSP-4 expression is associated with pro-inflammatory macrophages, we hypothesized that the regulation of macrophage number in peritonitis model may be associated with their ability to differentiate into the pro-inflammatory phenotype and to commit to apoptosis, a process closely associated with pro-inflammatory differentiation. Pro-inflammatory macrophages produce and release inflammatory stimuli, and this release is associated with their transition to apoptosis [46–53]. Both our *in vivo* and *in vitro* results suggested that TSP4 is needed for completion of pro-inflammatory differentiation of macrophages, their transition into apoptosis and their release of pro-inflammatory signals. This sequence of events was observed using cultured RAW264.7 and BMDM. TSP-4 KO reduced caspase 3 gene expression and activity of caspase 3/caspase 7, as well as the ratio of expressed Bax/Bcl2 in BMDM in response to LPS stimulation, confirming that TSP-4 promotes apoptosis in pro-inflammatory macrophages. P387 TSP-4 BMDM from P387-TSP4-KI mice had increased expression of apoptotic markers compared to BMDM isolated from WT mice, consistent with higher activity of this TSP-4 variant in many cellular effects and interactions that we previously reported.

TSP4 increased accumulation of macrophages in peritoneal tissues, presumably by increasing their adhesive properties and reducing the macrophage migration as was demonstrated in cultured cells. LPS injection significantly increased the number of macrophages recovered in the lavage of the peritoneal cavity of WT mice and *Thbs4*^−/−^ mice, but, in P387-TSP4-KI mice, the number of recovered macrophages was not increased by LPS compared to the saline control. Peritoneal macrophages in the lavage produced very little TSP4, while the levels of TSP4 mRNA and protein were increased in peritoneal tissue as was detected by QRT-PCR and immunofluorescence. TSP4 production was associated with CD68 positive cells, macrophages, in the tissue. The production of TSP4 by the blood cells has not been previously reported, and the ability of macrophages to produce TSP4 protein has not been recognized. However, as we report here, we identified several proinflammatory agonists that stimulate macrophages to produce TSP4 both in cell culture and *in vivo*.

P387-TSP4 is a SNP variant of TSP4 that is carried by >30% of North American population [18]. In our previous studies of this TSP4 variant *in vitro* in cell culture, and *in vivo* in a mouse model, we found that P387-TSP4 was more active in all cellular effects and interactions tested and was is less susceptible to proteolytic degradation [5,14,24,25,35–37,45]. Thus, the lack of increased macrophage egress into the peritoneal cavity of P387-TSP4-KI mice was surprising and suggested that P387-TSP4 prevented macrophage egress from tissues leading to increased accumulation in peritoneal tissues due to its higher adhesive properties than the WT, A387-TSP4 form.

TSP4 supports the adhesion of leukocytes [14,35], and macrophages specifically [14]. We have performed adhesion and migration assays with macrophage-like cells RAW264.7 and demonstrated that both A387-and P387-TSP4 support the adhesion of macrophages and reduce their migratory activity, but, as was expected based on our previous publications [24,35–37], P387-TSP4 is more effective in these functions.

Although the accumulation of macrophages in peritoneal tissue was increased in response to LPS in all three mouse strains, WT expressing the A387-TSP4 isoform, *Thbs4*^−/−^, and P387-TSP4-KI, the relative increase in macrophage accumulation (in comparison to mice receiving PBS) was surprisingly higher in *Thbs4*^−/−^ mice and significantly lower in P387-TSP4-Ki mice. Since the number of monocytes in blood was not significantly different between the three genotypes, and the number of macrophages in peritoneal cavity could not account for any differences in macrophage accumulation in tissues, these results suggested that TSP4 levels might affect survival of macrophages in tissues. In addition to the effect of TSP4 on CD68 levels, CD38 levels also followed the trend, suggesting that TSP4 acts on pro-inflammatory macrophages rather than tissue repair macrophages. Similar to other TSP4 functions, the P387 variant of TSP4 was more active in exerting these effects on macrophages, promoting inflammation, accumulation of macrophages in tissue, and their transition to apoptosis that allows macrophages to release pro-inflammatory stimuli and sustain a pro-inflammatory environment.

The association of TSP4 with tissue inflammation and pro-inflammatory differentiation of macrophages was observed *in vivo*, in peritonitis model, as well as in cultured macrophages. TSP-4 carries out a dual role in inflammation by facilitating macrophage adhesion and promoting the pro-apoptotic response of macrophages at sites of tissue inflammation. This association suggests a new and unanticipated role for TSP-4 in inflammation: TSP4 and its P387 variant are produced by macrophages in response to inflammatory stimuli and regulate the accumulation of macrophages in tissues and their pro-inflammatory functions.

## Abbreviations

bFGF: basic fibroblast growth factor
DMSO: dimethyl sulfoxide
ECM: extracellular matrix
MPM: mouse peritoneal macrophages
IP: intra-peritoneal
*Thbs4*−/−: thrombospondin-4 gene knock-out
WT: wild type
P387-TSP4 KI: 387P *TSP-4* knock-in mice
OCT: Optimum Cutting Temperature
vWF: von Willebrand factor
α-SMA: alpha-smooth muscle actin
Egr2: Early Growth Response 2
PBS: Phosphate Buffer saline
CHX: Cycloheximide
DMEM: Dulbecco’s Modified Eagle Medium
BMDM: Bone Marrow Derived Macrophage, LPS, lipopolysaccharide
GM-CSF: Granulocyte-Macrophage Colony-stimulating Factor
M-CSF: Macrophage Colony-stimulating Factor
INFγ: Interferon Gamma

## Acknowledgments

This work has been supported by R01 HL117216 (E.F.P. and O.S.A.) and R01 CA177771 (O.S.A.)

