## Supplemental data for "Effect of Thrombospondin-4 on Pro-inflammatory Phenotype Differentiation and Apoptosis in Macrophages"

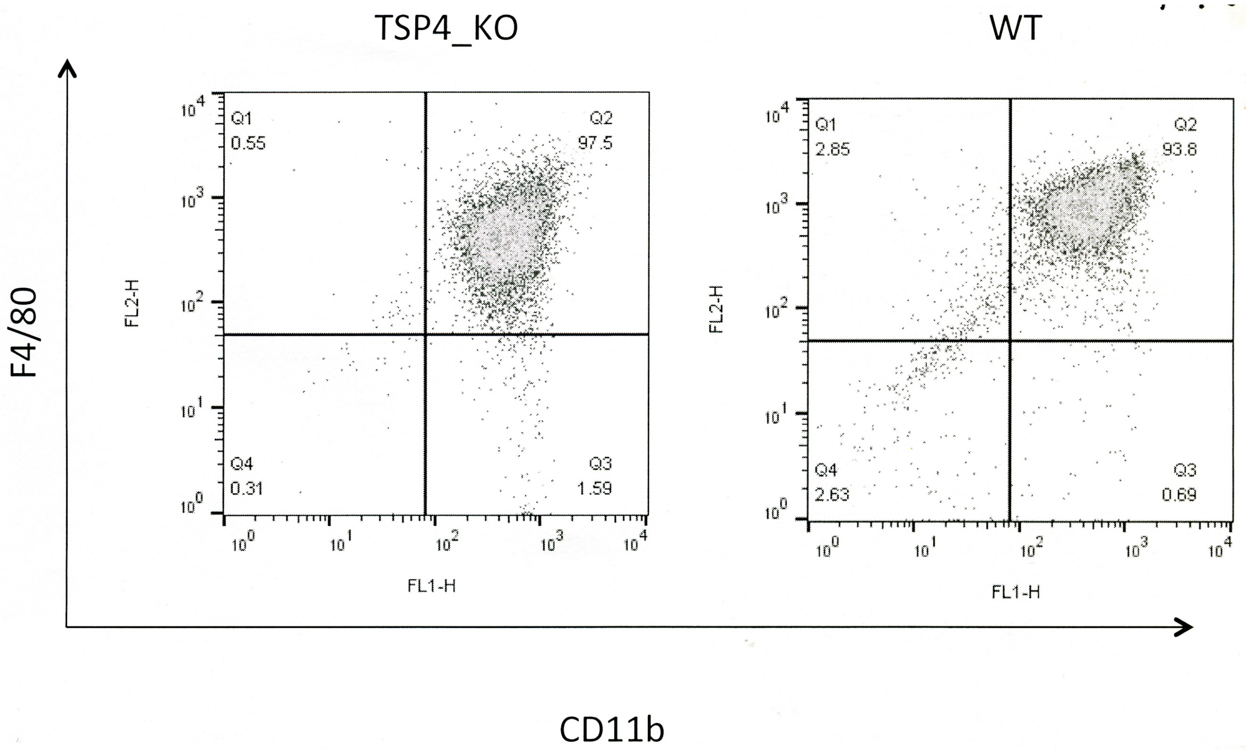


**Suppl. Figure 1. Purity of mouse bone-marrow-derived macrophages (BMDM).** BMDM were analyzed by flow cytometry using anti-CD11b and anti-F4/80 antibodies after differentiation in macrophage differentiation medium. 94 – 98 % of cultured cells were positive for both markers.

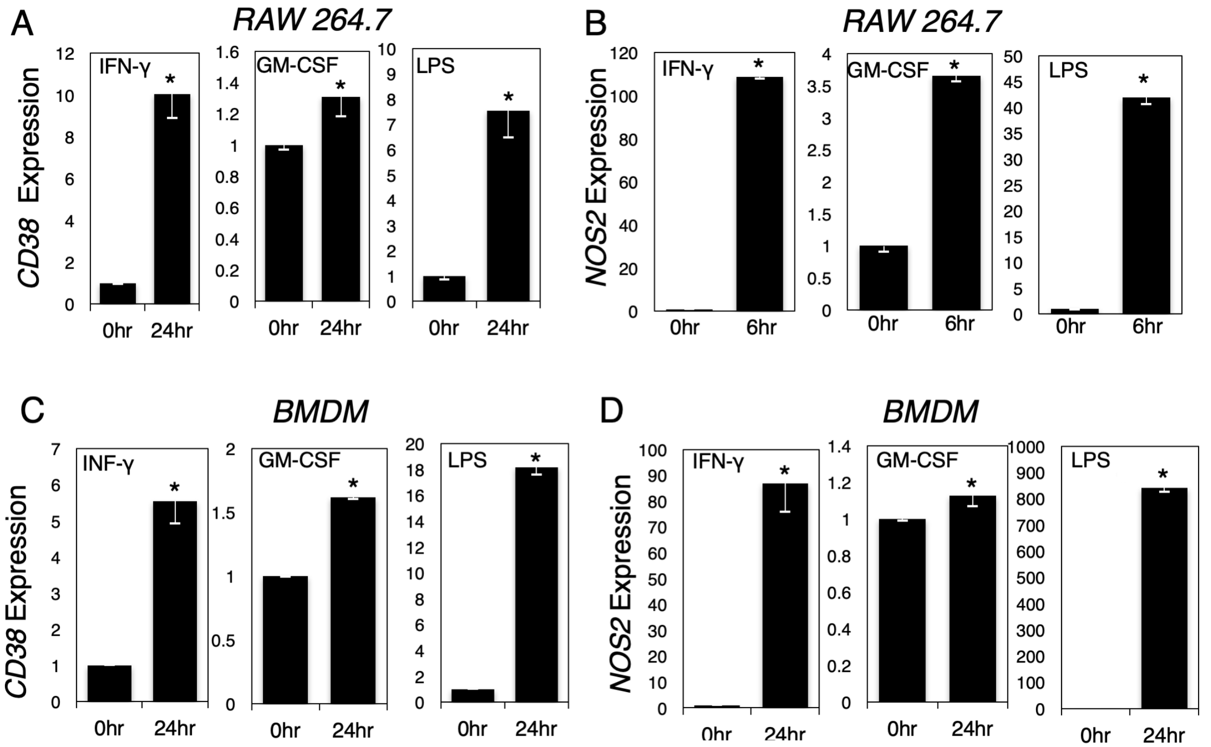


**Suppl. Figure 2. Expression of markers of pro-inflammatory macrophages in response to stimulation with pro-inflammatory stimuli.** Cultured RAW264.7 (A, B) and BMDM (C, D) were stimulated as described in Methods, and mRNA of the markers CD38 and NOS2 was analyzed by QRT-PCR. Fold increase over unstimulated cells, C = control, no stimulation; n = 3; *p < 0.05.

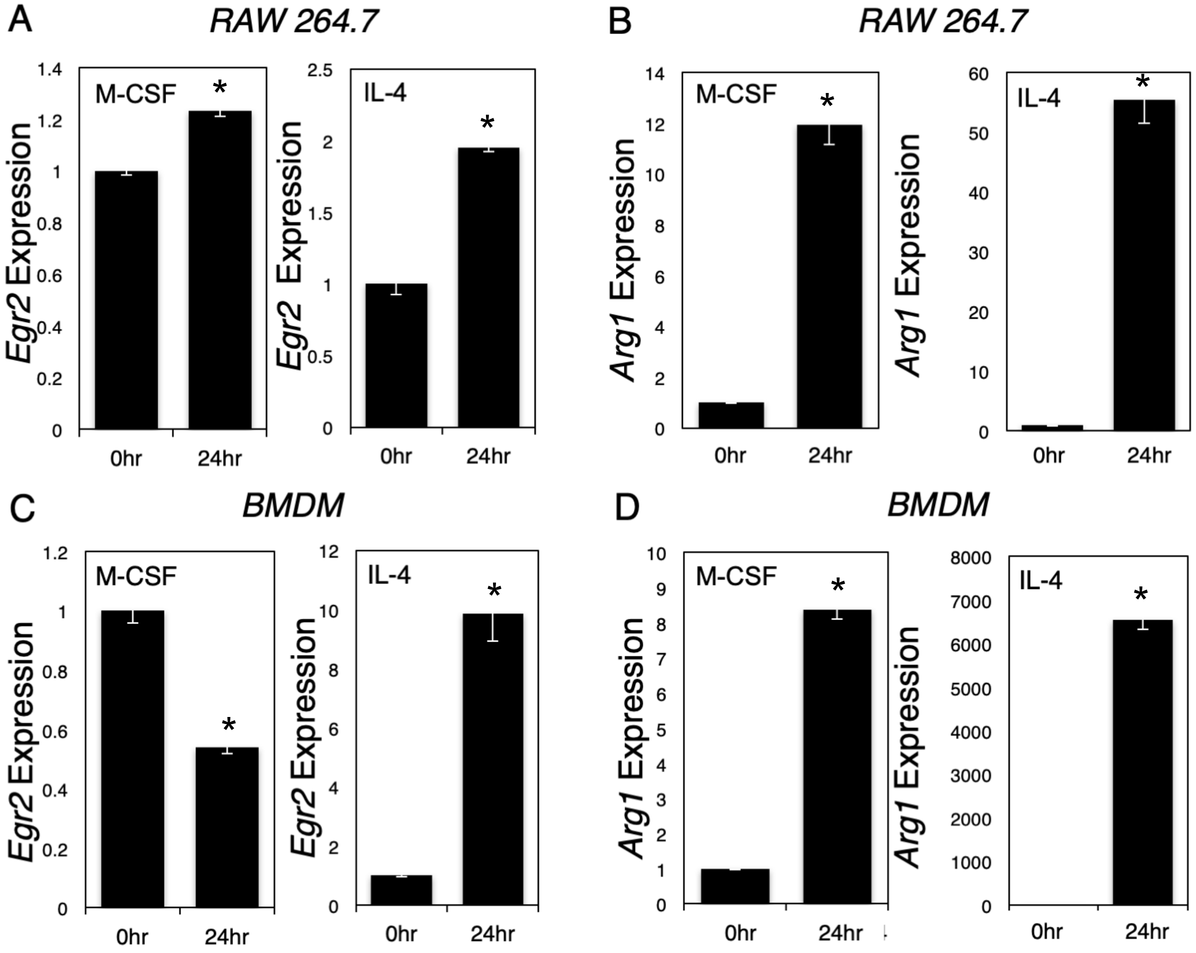


**Suppl. Figure 3. Expression of markers of tissue repair macrophages in response to stimulation with anti-inflammatory stimuli.** Cultured RAW264.7 (A, B) and BMDM (C, D) were stimulated as described in Methods, and mRNA of the markers Egr2 (A, C) and Arg1 (B, D) was analyzed by QRT-PCR. Fold increase over unstimulated cells, C = control, no stimulation; n = 3; *$p < 0.05$.

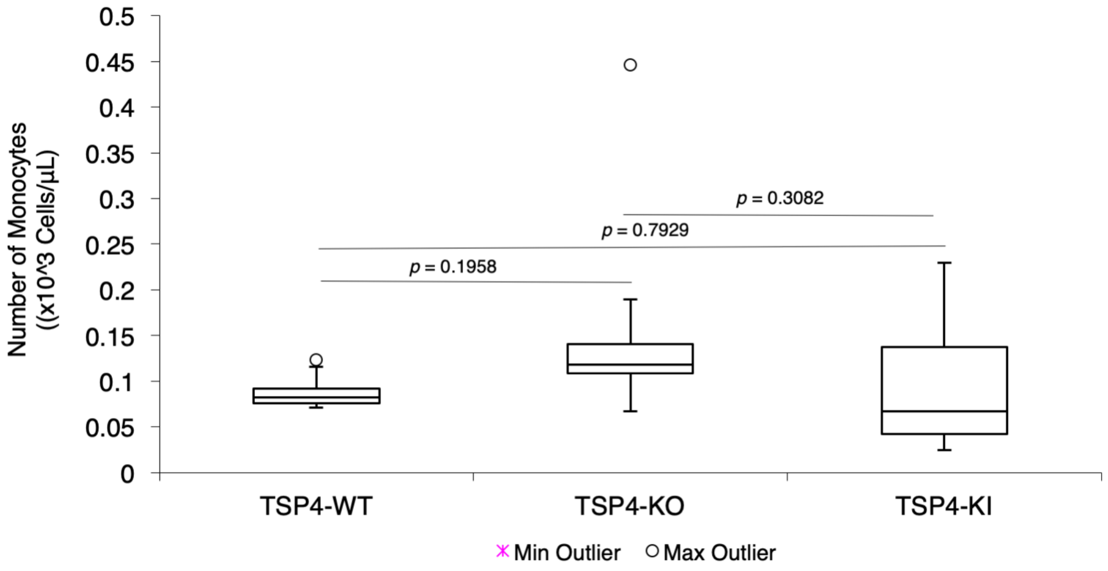


**Suppl. Figure 4. Number of blood monocytes in TSP4-WT (expressing A387-TSP4), Thbs4-/- (TSP4-KO), and P387-TSP4-KI (TSP4-KI) mice.**
